# Modeling the dynamics of antibody–target binding in living tumors

**DOI:** 10.1101/2020.05.12.090241

**Authors:** Yu Tang, Yanguang Cao

**Author notes:** Corresponding author: Yanguang Cao, UNC Eshelman School of Pharmacy, UNC at Chapel Hill.

## Abstract

Antibodies have become an attractive class of therapeutic agents for solid tumors, mainly because of their high target selectivity and affinity. The target binding properties of antibodies are critical for their efficacy and toxicity. Our lab has developed a bioluminescence resonance energy transfer (BRET) imaging approach that directly supports measurement of the binding dynamics between antibodies and their targets in the native tumor environment. In the present study, we have developed a spatially resolved computational model to analyze the longitudinal BRET imaging of antibody–target binding and to explore the mechanisms of biphasic binding dynamics between cetuximab and its target, the epidermal growth factor receptor (EGFR). The model suggested that cetuximab bound differently to EGFR in the stroma-rich area than in stroma-poor regions, and this difference was confirmed by immunofluorescence staining. Compared to the binding in vitro, cetuximab bound to EGFR to a “slower-but-tighter” degree in the living tumors. These findings have provided spatially resolved characterizations of antibody–target binding in living tumors and have yielded many mechanistic insights into the factors that affect antibody interactions with its targets.

## Introduction

The therapeutic antibody is an important class of therapeutics for treating solid tumors. More than 30 therapeutic antibodies have been approved for treating tumors at various stages^1, 2^. These broad applications of therapeutic antibodies in solid tumors are due, at least in part, to their high target binding selectivity and affinity when compared with traditional cytotoxic agents. Once bound to their targets, therapeutic antibodies eradicate tumor cells mainly by three mechanisms: blocking the pathogenic ligand–receptor interactions, triggering cell apoptosis pathways, or activating host effector functions^3^. The mechanisms of action are not exclusive but usually differ depending on the design of the different classes of antibodies.

Regardless of the mode of antibody action, antibody–target engagement is the first and most critical step for antibody efficacy. The patterns of target binding are often associated with the cellular response of the target cells and with treatment efficacy. Many studies have revealed that tumor cells can receive information by altering the temporal behavior (dynamics) of their signaling molecules^4^. A classic example of this behavior is the extracellular signal-regulated kinase pathway for epidermal growth factor receptor (EGFR). Transient activation (or blocking) of EGFR is associated with tumor cell proliferation, whereas sustained activation can lead to cell differentiation^5^. In addition, once antibodies have bound to their target cells, they can direct effector cells to elicit antibody–dependent cellular cytotoxicity (ADCC). Thus, the residence time of the antibody–target complex on tumor cells (determined by the off-rate) becomes critical for increasing lipid raft formation and the probability of ADCC^6^. Many tumor cells can initiate fast endocytosis upon antibody binding, leading to resistance to antibody attack^7^. Therefore, different target binding patterns can lead to distinct cellular reactions and treatment responses.

The target binding affinity is often assessed in vitro, using either surface plasmon resonance (SPR) or ligand competition assays. In SPR analysis, the antibody binds to target molecules that are immobilized on the sensing layer. Binding leads to changes in conformation and the angle of reflectivity, from which the association (k_on_) and dissociation (k_off_) can be quantified^8^. As in other routinely applied technologies that measure binding dynamics, the k_on_ and k_off_ that are determined by SPR merely reflect antibody–target interactions at the molecular level. These techniques are valuable for antibody screening, but they are not relevant to binding under physiological conditions. The target binding properties in living systems remain largely uncharacterized.

Tumor tissues are known to be very heterogeneous, both between and within tumors. In addition to complex tumor genotypes, morphological and phenotypic features can differ, even within the same tumor. The stromal environment where each tumor cell resides largely shapes its phenotypic properties^9^. However, how these stromal components can influence the binding dynamics between an antibody and its target cell remains largely undefined. Unlike *in vitro* assay systems, where all targets are freely accessible, tumors present many physical barriers that influence the diffusion of antibodies, as well as their interactions with the target^10^. Previous studies have shown that antibodies are unable to freely reach their target or cannot drift away after dissociating from the target in the presence of spatial obstacles^11^.The resulting shifts in binding dynamics within living tumors can reduce the cellular response or even lead to treatment failure.

We have developed a bioluminescence resonance energy transfer (BRET) imaging system that can directly monitor the antibody–target binding dynamics in living systems^12^. This imaging system leverages a high signal-to-noise ratio and stringent energy donor-acceptor distance to provide specific measurements of antibody–target binding dynamics in a selective and temporal fashion. It is a minimally invasive system, so it enables longitudinal monitoring of in vivo antibody–target interactions. We have previously used this approach to demonstrate that cetuximab binds to its target in a biphasic and dose-shifted manner. In the present study, we have developed spatially resolved computational models for analysis of the longitudinal imaging data of antibody–target binding in living tumors and we have compared their binding dynamics in spatially distinct tumor areas. With these models, we have assessed possible mechanisms that could explain the biphasic features of cetuximab–EGFR binding in a xenograft tumor. The results of this study have provided many insights into the dynamic features of antibody–target binding in living tumors and the stroma factors that potentially influence those dynamics.

## Methods

### Study Design

Our lab has developed a BRET approach to support the investigations of antibody–target binding dynamics in the native tumor environment. Specifically, a small but bright luciferase, NanoLuc, was fused to the extracellular domain of EGFR to serve as the energy donor in the BRET pair^12^. An anti-EGFR antibody, cetuximab, was labeled with DY605, a fluorophore with an emission wavelength at 625 nm, to serve as the energy acceptor. Prior to the binding between DY605-labeled cetuximab (DY605-CTX) and the NanoLuc-fused EGFR (NLuc-EGFR), the distance between NanoLuc and DY605 was too great to trigger BRET, and only the bioluminescence emission at 460 nm for NanoLuc was observed. However, binding of DY605-CTX to NLuc-EGFR increased the proximity between NanoLuc and DY605 and allowed the transfer of bioluminescence energy to DY605 and the emission of fluorescence signals at 625 nm.

The experimental design is shown in **Fig. 1**. In total, 20 nude mice were inoculated with NLuc-EGFR-expressing HEK293 cells; and the BRET imaging study was performed when the tumor sizes had reached 500 mm^3^. The imaging study was initiated by injecting the DY605-labeled cetuximab via the tail vein of the xenograft mice at three doses: 1.0, 8.5, and 50 mg/kg (n = 5 mice/dose group), or DY605-labeled human IgG (n = 5/dose group). Blood samples (30 μL) were collected at the designated times for PK assessment. The plasma concentrations of cetuximab were quantified based on fluorescent intensities. Images at both 460 nm and 625 nm were acquired using an IVIS Kinetic optical imaging system (Caliper Life Sciences, Alameda, CA, USA) upon intraperitoneal (i.v., 0.25 mg/kg) administration of substrate (0.25 mg/kg). The fluorescence intensity was determined to quantify the concentrations of the antibody–target complex, and the receptor occupancy (RO) was derived using the following equation.

**Figure 1.**
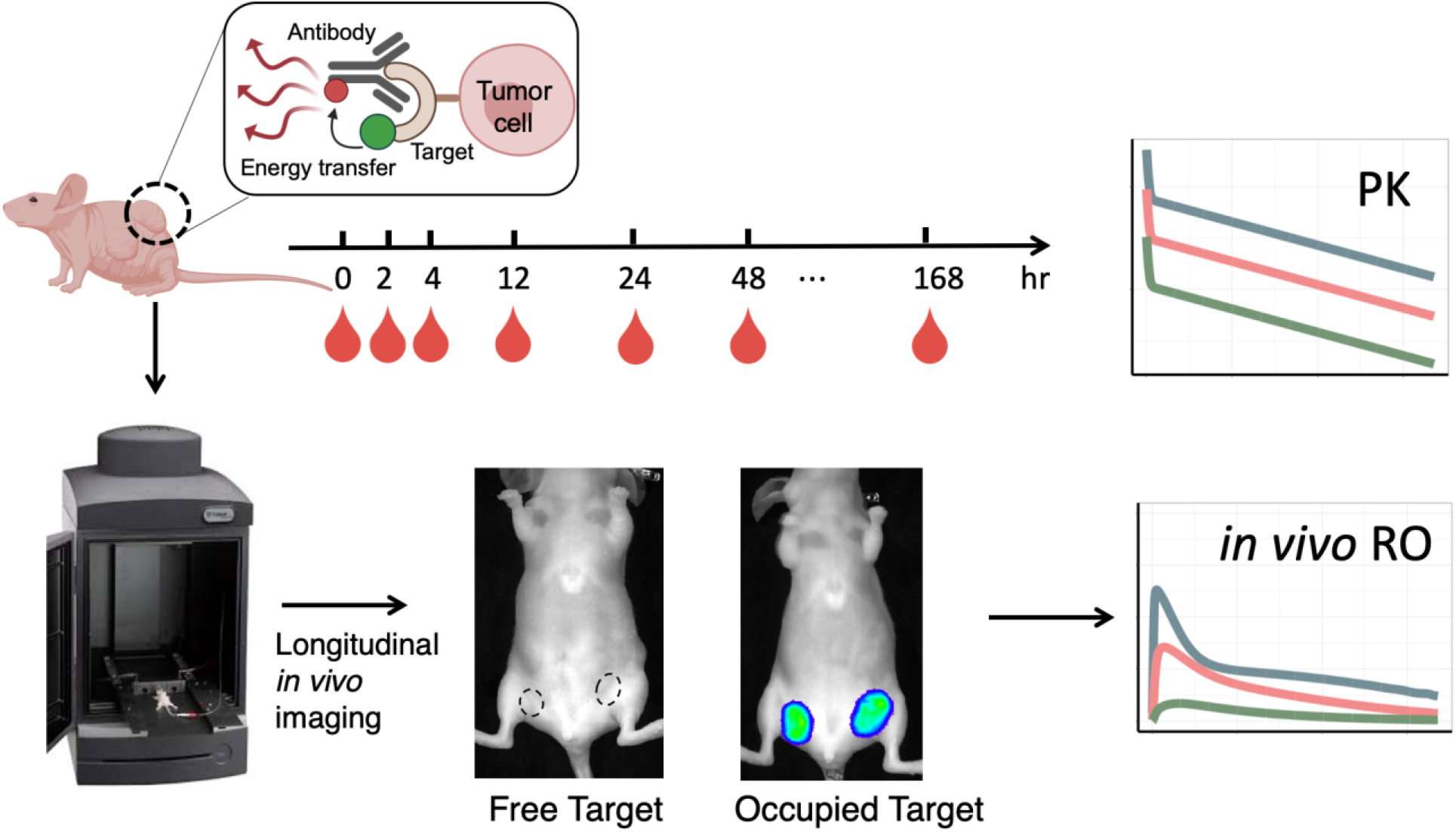
The scheme of the experimental design of the Bioluminescence Resonance Energy Transfer (BRET) study. A small, but bright, luciferase, NanoLuc, was fused to the extracellular domain of EGFR to serve as the energy donor in the BRET pair (Nluc-EGFR). The anti-EGFR antibody cetuximab was labeled with DY605 as the energy acceptor (DY605-CTX). Twenty nude mice were inoculated with NLuc-EGFR-expressing HEK293 cells; and the BRET imaging study was performed after xenograft tumor sizes reached 500 mm^3^. DY605-CTX at three doses (1.0, 8.5, and 50 mg/kg) or DY605-human IgG was injected via tail vein (n = 5/dose group). Blood samples (30 μL) were collected at designated times for pharmacokinetics assessment. The plasma concentrations of DY605-CTX were quantified based on fluorescence intensities. Images were acquired using an IVIS Kinetic optical imaging system upon administration of the NanoLuc substrate furimazine (i.v., 0.25 mg/kg). The fluorescence intensity was determined to quantify the concentrations of the antibody–target complex and to derive the receptor occupancy (RO). The tumors were collected at the end of the study (around 192 h) and snap-frozen in liquid nitrogen.

The tumors were collected at the end of the study (at approximately 192 h) and snap-frozen in liquid nitrogen.

### Plasma PK model

The antibody plasma PK was described using a two-compartment model with a linear tissue distribution (CL_D_) and a linear systemic clearance (CL_P_) (**Fig. 2**). The PK data in three dose groups (1.0, 8.5, and 50 mg/kg) were analyzed simultaneously using the PK model, using a naïve pooled-data (NPD) approach. The volume of plasma (V_plasma_) was set to 0.001 L for 20 g mice^13^.

**Figure 2.**
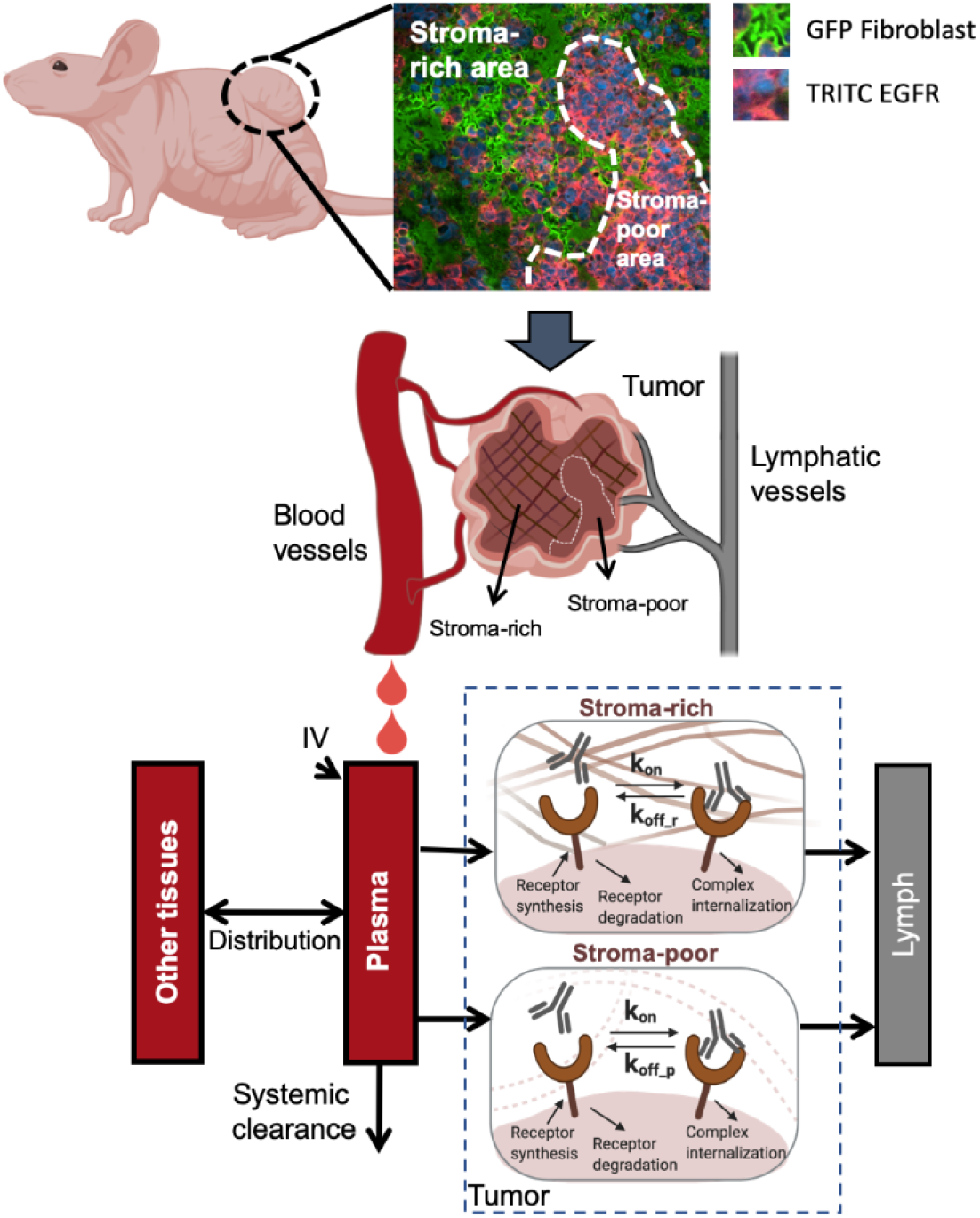
The spatially resolved computational model describing the antibody–antigen binding kinetics in xenografts. The antibody plasma pharmacokinetics^14^ were described using a two-compartment model with a linear tissue distribution and a linear systemic clearance. The solid tumors were conceptually dissected into two anatomical compartments—stroma-rich and stroma-poor areas—to account for the spatial heterogeneity. The stroma-poor compartment described the area where tumor cells grow without any spatial restriction by stromal cells, whereas the stroma-rich tumor compartment represented the area where tumor cells grow in the presence of dense tumor-associated stromal cells (e.g., fibroblasts). Antibodies were assumed to extravasate from tumor blood vessels into the interstitial space and leave the interstitial space via lymphatic vessels. In both tumor compartments, the free receptors were synthesized and degraded on the tumor cells. The antibody–receptor complexes were cleared by internalization. The free antibodies bound to free receptors at a rate of k_on_. The antibodies dissociated from receptors at a rate of k_off_r_ in the stroma-rich compartment and a rate of k_off_p_ in the stroma-poor compartment.

### Modeling antibody–target binding dynamics in tumors

The dynamics of antibody–target binding in solid tumors were further characterized to obtain mechanistic insights by implementing a sequential modeling strategy. Here, the PK model was first optimized and then fixed during the second step to explore the antibody–target binding dynamics.

The solid tumors were conceptually dissected into two anatomical compartments: a stroma-rich and a stroma-poor area, to account for the spatial heterogeneity (**Fig 2**). The stromal-rich compartment described the area where tumor cells grew relatively quickly, without any spatial restriction by stromal cells. By contrast, the stroma-poor tumor compartment represented the area where tumor cells grew in the presence of dense tumor-associated stromal cells (e.g., fibroblasts). The relative volume and blood flow in the two tumor areas were evaluated as model parameters.

The extravasation of the antibody from the tumor blood vessels to the interstitial space was assumed to be dominated by convection and was described by a vascular reflection coefficient (σ_v_) and the convective lymph flow into either the stroma-rich area (L_r_) or the stroma-poor area (L_p_)^7^. The value of σ_v_ was set at 0.78, a value reported for subcutaneous xenograft models^13^. The values of L_p_ and L_r_ were functions of the tumor blood flow (TBF)^14^, the total tumor volume (V_tumor_), and the relative fraction between the two tumor areas (f_t_), as described in the following equations:

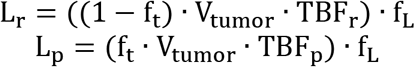

where TBF_p_ and TBF_r_ describe the tumor blood flows in two tumor areas. The value of f_L_ was set to 0.2%^15^. The total tumor volumes (V_tumor_) were measured using a caliper.

The spaces for antibody distribution and for antibody–target interaction in both tumor compartments were set to a fraction (f_av_) of the total interstitial space.

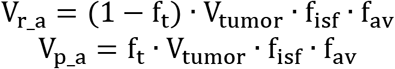

The available fraction f_av_ was set at 27.5% ^13, 16^. Notably, the antibody–antigen bindings occurred in the same space that was accessible to antibodies (V_r_a_ and V_p_a_). The target (i.e., EGFR) was assumed to be synthesized by tumor cells at a zero-order rate constant (k_syn_) and to be endocytosed at a first-order rate constant (k_deg_). The cetuximab-EGFR complex was assumed to be internalized by the tumor cells at a first-order rate constant (k_int_). All parameters regarding the target protein (k_syn_, k_deg_, and k_int_) were assumed to be conserved inside the tumors. The association rate between the antibody and target is denoted as k_on_ and the dissociation rate constant is k_off_.

We used the developed modeling framework primarily to investigate two competing hypotheses: (1) the antibodies bind to the targets differentially across two tumor areas (the heterogeneous binding model, HBM) and (2) the antibodies are distributed differentially into two tumor areas, but with the same binding profile (the heterogeneous distribution model, HDM). The differential equations for both models are provided in the Supplementary materials. We optimized both models against the data and evaluated which model made more consistent predictions to the observed RO data. The selection of the most suitable model and the parameter estimates were confirmed by visual inspection, by the observed versus predicted plot, by the predicted versus residual plot, by the CV% of estimated parameters, and by the physiological plausibility of the estimated parameters.

### Immunofluorescence (IF) staining

We assessed the spatial distributions of the antibody in tumors by IF staining after the imaging study. The tumor samples were preserved and sliced in OCT medium (23-730-571, Fisher scientific, Waltham, MA, USA). The sliced tumor tissues were fixed in methanol/acetone (1:1) at 4°C. After blocking with phosphate buffered saline (PBS) containing 2% fetal bovine serum (FBS) (F2442, Millipore Sigma, Burlington, MA, USA; 10010023, Thermo, Waltham, MA, USA), the tumor slices were stained with anti-EGFR and anti-α-SMA antibodies. In brief, the slides were incubated with Alexa Fluor 555-conjugated primary rabbit anti-human EGFR antibodies (MA7-00308-A555, Thermo, Waltham, MA, USA, 1:1000 diluted in PBS) and primary mouse anti-mouse α-SMA antibodies (14-9760-82, Thermo, Waltham, MA, USA, 1:1000 diluted in PBS) at 4°C overnight and then incubated with Alexa Fluor 555-conjugated goat anti-mouse antibodies (A-11001, Thermo, Waltham, MA, USA, 1:1000 diluted in PBS) at room temperature for 1 h. The immunofluorescence images were acquired with a Live Cell Imaging Microscope (Nikon, Melville, NY, USA).

### Model simulation

The developed model was applied to simulate the profiles of antibody–target binding dynamics and the resultant RO at different conditions. The concentrations of free antibodies, free targets, and antibody–target complexes in both tumor areas were simulated and compared. In addition, the SPR-measured binding parameters were applied to replace the optimized parameters to allow an examination of differences in antibody–EGFR binding dynamics in the living tumors versus in vitro binding in buffers.

## Results

### Plasma PK and antibody–target binding dynamics in tumors

In this study, DY605-cetuximab showed bi-exponential and linear PK profiles^12^, as the area under the curve (AUC) and the peak plasma concentrations increased proportionally to the doses. A temporal shift was observed from the antibody plasma PK to the ROs in the tumors. The tumor ROs peaked at approximately 4 h post-dosing, which was consistent across doses, suggesting that the extravasation of DY605-cetuximab into tumors is a slow and linear process. The increase in the tumor ROs was less than dose proportional, indicating a nonlinear process was involved in the conversion of free antibodies in the plasma to bound antibodies in tumors. Notably, the target EGFR in the tumors was not saturated, even at a supra-therapeutic antibody dose (50 mg/kg), suggesting fractional target accessibility. Furthermore, the RO profiles declined in a biphasic manner and showed a shallow terminal declining phase, particularly at the two higher doses. Interestingly, the transition from the rapid to the slowly declining phases was not consistent across doses.

### The antibody–target binding profiles in tumors were well recapitulated by the HBM

The average plasma concentrations and tumor ROs were used for model competition. As shown in **Fig. 3A**, the two-compartment PK model adequately recapitulated the PK profiles at all doses. This confirms the linear PK properties of DY605-cetuximab in xenograft mice within the assessed dose range. The estimated PK parameters and CV% are shown in **Table 1**. The estimated systemic clearance and half-life were consistent with those of previous reports.

**Figure 3.**
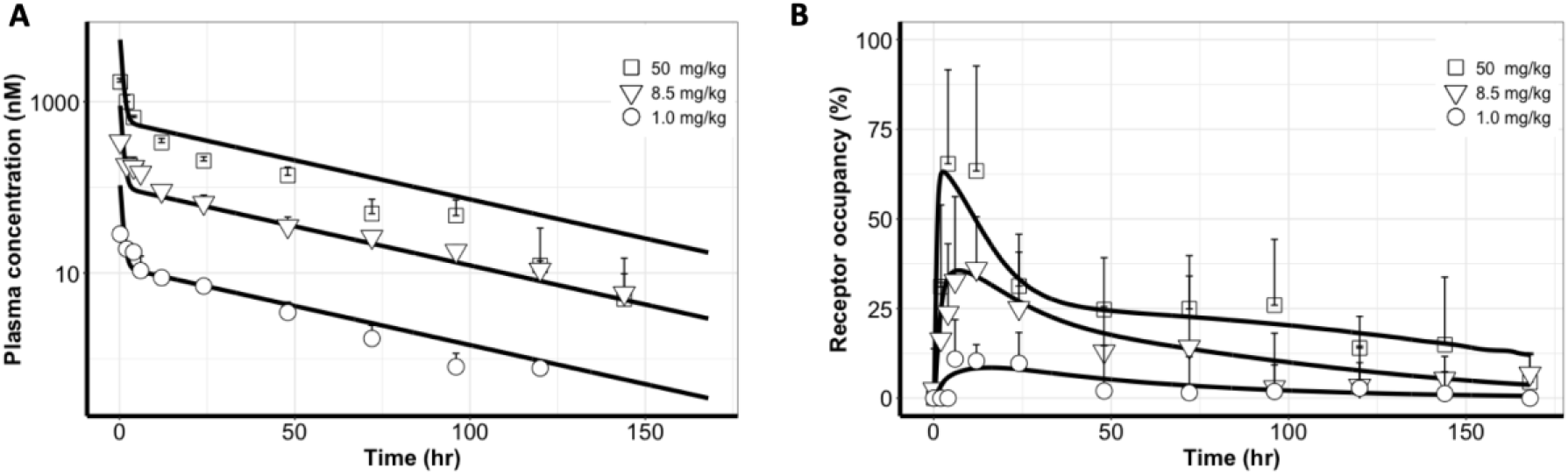
The profiles of antibody–target binding in tumors were well recapitulated by the heterogeneous binding model (HBM). **(A)** The two-compartment PK model adequately recapitulated the PK profiles at all doses. **(B)** The tumor receptor occupancy (RO) profiles were well characterized by the heterogeneous binding model (HBM) at three doses. Each data point represents the mean plasma concentration or mean RO. Error bars represent ±SD.

**Table 1.** Pharmacokinetics (PK) parameter estimations

| Parameter | Unit | Definition | Estimation (CV%) |
| --- | --- | --- | --- |
| CL<sub>D</sub> | L·hr<sup>-1</sup> | Tissue distribution flow | 0.0012 (15%) |
| CL<sub>p</sub> | L·hr<sup>-1</sup> | Cetuximab systemic clearance | 0.00021 (4.8%) |
| V<sub>other</sub> | L | Other tissue volume | 0.0076 (7.1%) |
| V<sub>plasma</sub> | L | Plasma volume | 0.001 (fixed) |

The tumor RO profiles were well characterized by the HBM at all three doses (**Fig. 3B and 4A**). The model suggested different antibody–target binding dynamics across two tumor spatial areas. The optimized parameters are shown in **Table 2**. The association rates (i.e., k_on_) between cetuximab and its target EGFR in both tumor compartments were estimated to be close and were therefore considered a shared parameter in the two tumor compartments. The estimate of k_on_ was 0.03 nM^-1^·h^-1^, a value approximately 1% of the rate measured using SPR^17^, indicating the impact of physical barriers on the association rate in the living tumors compared to the in vitro buffer conditions. Interestingly, the complex dissociation rate (i.e., k_off_) was markedly different between the two tumor areas. The optimized k_off_ was 0.61 h^-1^ in the stroma-poor tumor area, which is close to the SPR measured values. However, the complex dissociation rate was estimated as much slower (k_off_r_ = 0.0017 h^-1^) in the stroma-rich tumor area^17^.

**Figure 4.**
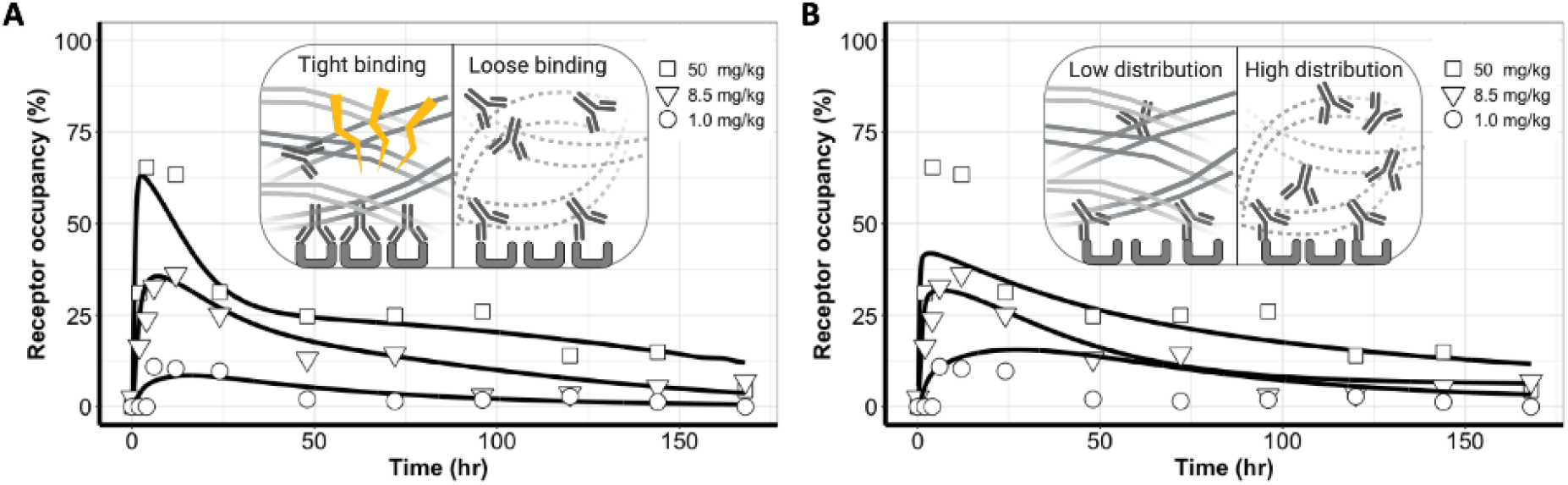
Uneven distributions of antibodies in tumors could not sufficiently explain the observed binding dynamic features in comparison to the heterogeneous binding patterns. **(A)** The heterogeneous binding model (HBM) well-captured the antibody–target binding kinetics, whereas **(B)** The heterogeneous distribution model (HDM) failed to capture the receptor occupancy (RO) profiles across three doses, and a clear model misspecification was indicated.

**Table 2.** Heterogeneous binding model (TBM) parameter estimations

| Parameter | Unit | Definition | Estimation (CV%) |
| --- | --- | --- | --- |
| $k_{deg}$ | $hr^{-1}$ | EGFR degradation rate | 0.013 (42%) |
| $R_0$ | nM | EGFR initial concentration in tumor stroma-rich and stroma-poor space | 0.0020 (73%) |
| $k_{on}$ | $nM^{-1} \cdot hr^{-1}$ | Cetuximab-EGFR association rate | 0.030 (53%) |
| $k_{off\_p}$ | $hr^{-1}$ | Cetuximab-EGFR dissociation rate in stroma-poor regions | 0.61 (55%) |
| $k_{off\_r}$ | $hr^{-1}$ | Cetuximab-EGFR dissociation rate in stroma-rich regions | 0.0017 (54%) |
| $f_t$ | | Ratio of tumor stroma-poor volume over total space | 0.89 (5.8%) |
| TBF | $hr^{-1}$ | Tumor Blood Flow per 1 L tumor | 9.2 (21%) |
| $k_{int}$ | $hr^{-1}$ | Cetuximab-EGFR internalization rate | 0.14 (26%) |

Unfortunately, as shown in **Fig. 4B**, the HDM failed to capture the RO profiles across the three doses and clear model misspecification was indicated. Even with a sharp distribution gradient across tumor areas, the model could not provide a good prediction of the biphasic dynamic feature in the RO profiles. The RO peaks at 50 mg/kg were drastically under-predicted, while the RO values at 1.0 mg/kg were over-predicted. The poor performance of HDM indicated that heterogeneous antibody distribution could not be the primary mechanism for the biphasic declining feature of antibody–target binding in tumors. The parameter estimations of HDM are presented in **Supplementary Table 1**.

The different performance between HBM and HDM was also indicated by the Akaike information criterion (AIC). We concluded that the biphasic dynamic feature of RO was better explained by the different binding profiles than by the different distribution profiles in the two tumor compartments. Therefore, we selected the HBM as the model for further exploration.

### Cetuximab persisted longer in the stroma-rich area than in the stroma-poor area

At the end of the imaging study, we sectioned the tumors and stained the EGFR-positive tumor cells, tumor-associated fibroblasts, and cetuximab to evaluate the residual antibodies and their spatial distributions. **Fig. 5** shows representative images of the spatial distribution of the antibody (DY605-CTX), tumor-associated fibroblast (GFP Fibroblasts), and EGFR-positive tumor cells (TRITC EGFR). Area P represents the tumor area without many stroma cells and with evenly distributed tumor cells. Area R represents the stroma-rich area, where tumor cells were surrounded by tumor-associated fibroblasts. As shown in **Fig. 5**, by 192 h after antibody dosing, some antibodies were still present in the stroma-rich area, while detection of antibodies in the stroma-poor area was negligible. No residual antibodies were observed for non-specific IgG, suggesting that the residual antibodies were associated with Fab binding and not with nonspecific binding (**Supplementary Figure 1**).

**Figure 5.**
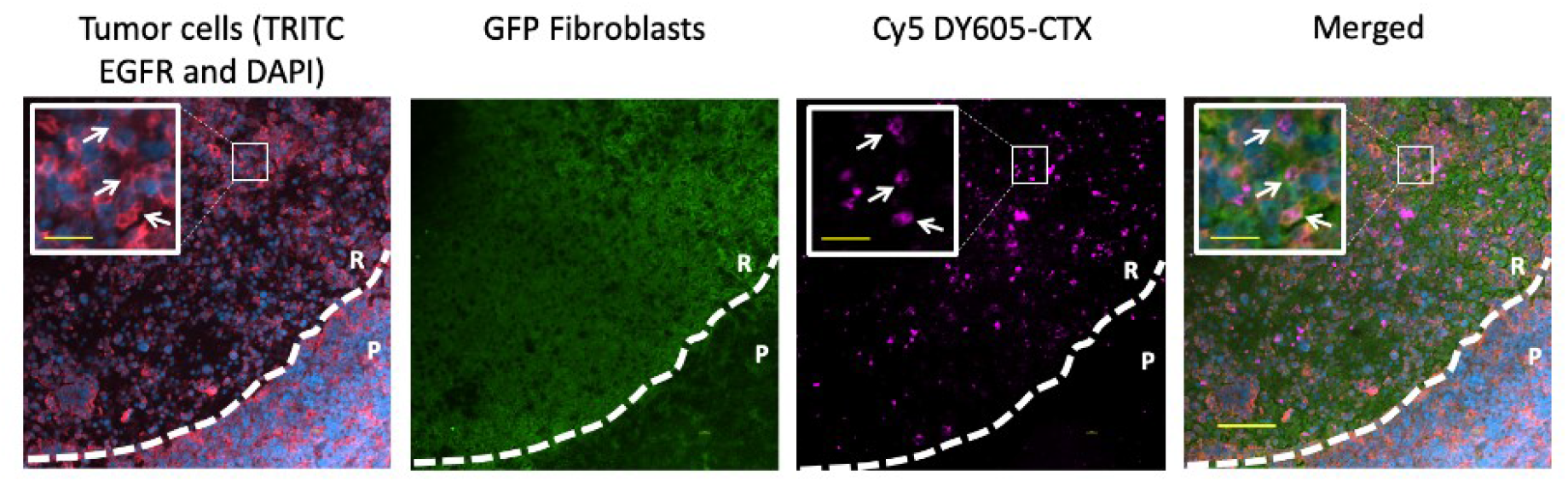
Cetuximab persisted longer in the stroma-rich area than in the stroma-poor area. A representative immunofluorescence (IF) image shows the histology of the tumor collected at the end of the bioluminescence resonance energy transfer (BRET) imaging study, revealing the spatial distribution of the antibody (Cy5 DY605-CTX), tumor-associated fibroblast (GFP Fibroblasts), and EGFR-positive tumor cells (TRITC EGFR and DAPI). Area P represents the tumor area without many stroma cells and with evenly distributed tumor cells. Area R represents the stroma-rich area, where tumor cells were surrounded by tumor-associated fibroblasts. As the white arrows indicate, residual antibodies are present at the tumor cell surfaces and largely overlap with EGFR staining, indicating that the detected Cy5 signals represent the bound DY605-CTX.

A close inspection of the spatial location indicated that residual antibodies had a very high co-localization with tumor cells (**Fig 5**). We observed that most of the residual antibodies in the stroma-rich area were retained on the surfaces of the tumor cells, likely as antibody–target complexes. This observation is consistent with our model predictions whereby antibodies would dissociate from the target much more slowly in the stroma-rich area. Only a small amount of bound antibodies was observed at the edge of Area P, and most antibodies had been degraded in the stroma-poor tumor area. In addition, the total EGFR was sparser in Area R than in Area P, suggesting a higher EGFR suppression in the stroma-rich tumor area. These findings agreed with the model simulations described in the next section.

### Cetuximab bound EGFR at a “slower-but-tighter” degree in living tumors than in the in vitro conditions

We further examined antibody binding dynamics in tumors in comparison to the binding in the in vitro condition, which is usually measured using SPR methods. We replaced the target binding parameters in the HBM with the SPR-measured values to predict the RO profiles at three doses, which were superimposed on the experimental observations. When k_on_ was set to an SPR-measured value in both tumor compartments, the model over-predicted the RO. Even though the biphasic feature on RO curves was predicted, almost no difference was detected in the predicted RO profiles across the three doses (**Fig. 6A**). With the SPR-measured k_off_ in both tumor compartments, the model under-predicted the RO data (**Fig. 6B**). The biphasic declining feature disappeared in this parameter setting. When both target binding parameters were set to SPR-derived values, the model also over-predicted ROs (**Fig. 6C**). Collectively, these findings confirmed a marked difference in antibody–target binding dynamics between the living tumors and the in vitro buffer systems.

**Figure 6.**
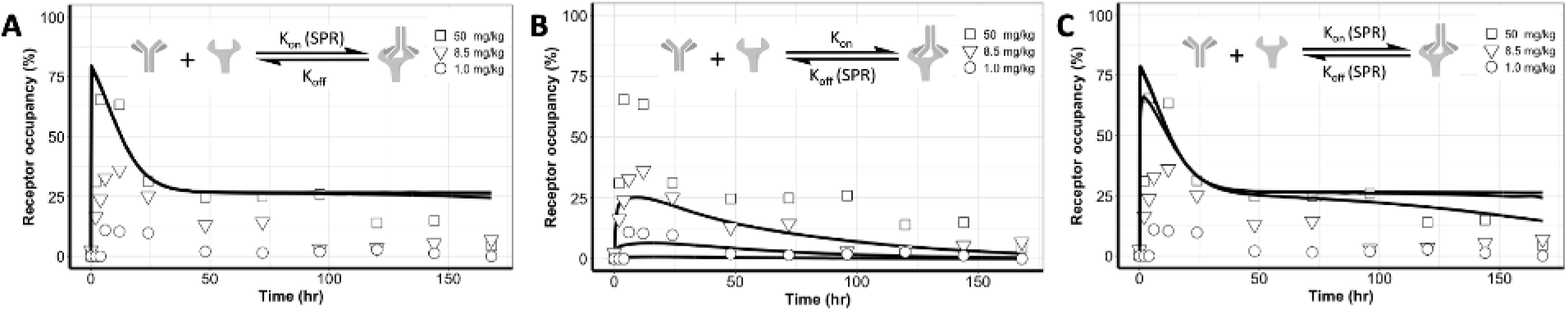
Cetuximab bound EGFR at a “slower-but-tighter” degree in the living tumors than in the in vitro conditions. The target binding parameters in the heterogeneous binding model (HBM) were replaced with the SPR-measured values to predict the receptor occupancy (RO) profiles at three doses; these are superimposed on the experimental observations. **(A)** When k_on_ was set to a SPR-measured value in both tumor compartments, the model over-predicted the RO. No difference was detected in the predicted RO profiles across the three doses. **(B)** When a SPR-measured k_off_ was used in both tumor compartments, the model under-predicted the RO data. **(C)** SPR-measured k_on_ and k_off_ values could not differentiate the RO profiles across the three doses.

### Cetuximab durably suppressed free EGFR but transiently formed antibody–target complexes in living tumors

We simulated free cetuximab, free EGFR, antibody–antigen complexes, and the RO in both tumor areas. The free cetuximab was similar in both tumor compartments at all doses (**Fig. 7A**), consistent with the model assumption. Free EGFR was rapidly suppressed, and the suppression lasted for over 150 h, particularly at 50 mg/kg (**Fig. 7B**). Cetuximab was predicted to have a stronger suppressive effect on free EGFR in the stroma-rich area. The magnitude and duration of EGFR suppression were both dose dependent, so a higher antibody dose gave a larger and longer suppression of the free target (**Fig. 7B**).

**Figure 7.**
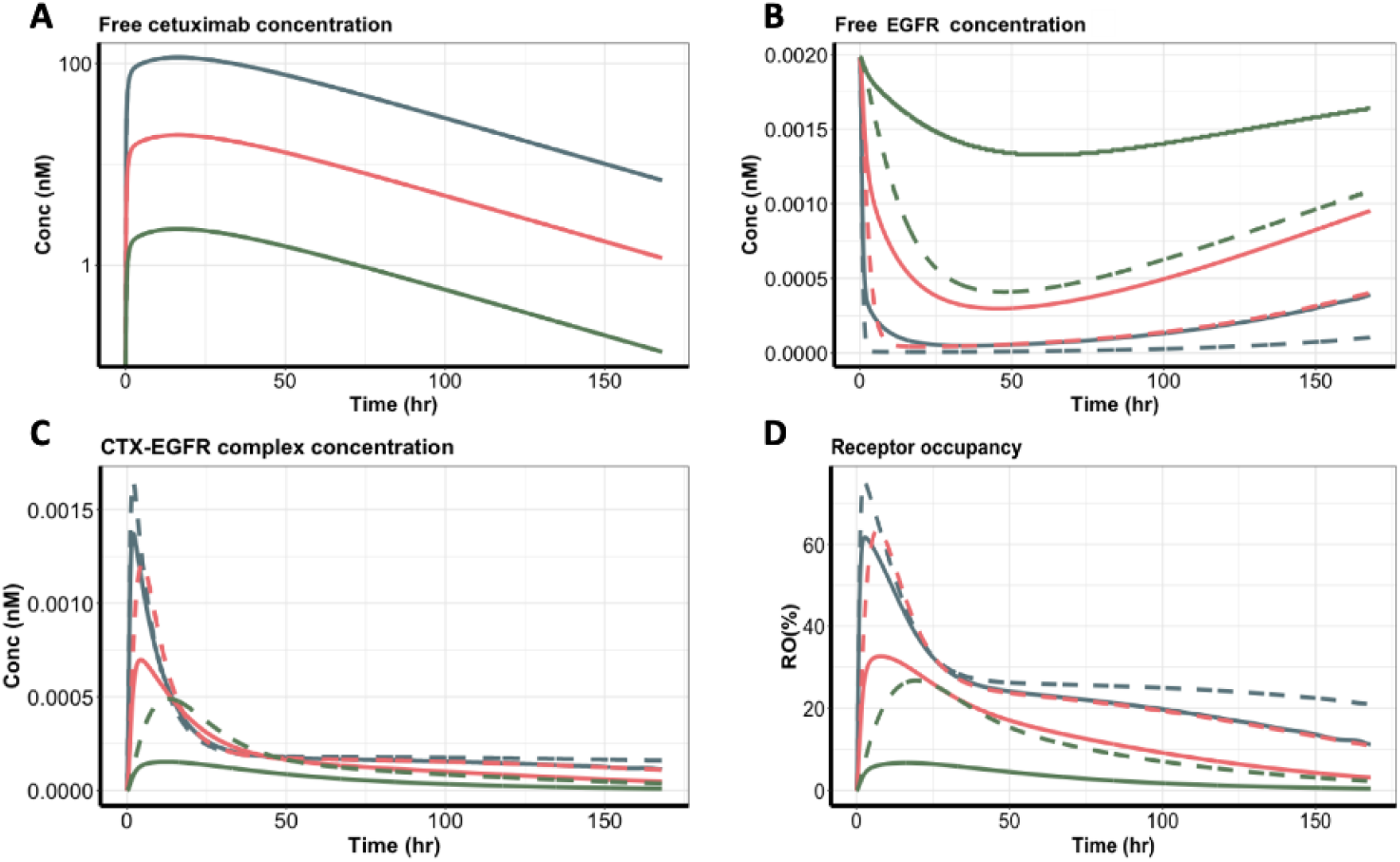
Cetuximab durably suppressed free EGFR, but the complex was formed transiently in tumors. **(A)** The free cetuximab concentration, **(B)** free EGFR concentration, **(C)** cetuximab-EGFR complex concentration, and **(D)** receptor occupancy in the two tumor compartments were simulated based on the heterogeneous binding model (HBM). Blue, red, and green lines represent the three cetuximab dose groups: 50, 8.5, and 1.0 mg/kg. Solid lines represent the tumor stroma-poor compartment, whereas the dashed lines represent the tumor stroma-rich compartment.

Compared to the durable target suppression, the formation of the antibody–target complex was quite transient (**Fig. 7C**). In both tumor compartments, the complex concentrations peaked at around 5 h after dosing, but the complex decayed shortly after the peaks and was subsequently maintained at relatively low levels for an extended period. A small difference was evident in complex concentrations across doses during the terminal phase. Despite the similar concentrations in the two tumor areas, the different binding properties of cetuximab meant that it showed relatively higher and more durable target properties in the stroma-rich than in the stroma-poor areas (**Fig. 7D)**. The difference between the two tumor areas became smaller as the dose increased.

## Discussion

Our understanding of antibody–target interactions, and particularly in the native physiological context, is still limited, mainly due to the lack of approaches to detect their binding dynamics with temporal and spatial resolution or specificity^18^. For example, immunohistochemistry (IHC) staining can quantify the spatial distribution but it often fails to incorporate the dynamic factors present in physiological situations^19^. Most in vivo imaging methods often cannot distinguish signals arising due to specific target engagement versus nonspecific signals^12^. We previously developed a BRET method that enables longitudinal monitoring of the binding dynamics between the antibody and its target in living tumors^12^. Using this method, we observed biphasic and dose-shifted binding dynamics between cetuximab and its target EGFR. In the present study, we developed a spatially resolved computational model to disentangle the dynamic binding patterns and evaluate the mechanisms.

Compared to the in vitro systems, an antibody in a living tumor could bind to its target to a “slower-and-tighter” degree. We observed that the antibody bound differently to its target in the stroma-rich areas than in the stroma-poor regions. Antibodies had a much slower dissociation rate in the stroma-rich areas, which was confirmed by immunofluorescence staining. The binding features in the stroma-rich tumor area were consistent with experimental observations that the stress stroma could restrict the diffusion of antibodies in the tumors^20^. This restricted diffusion could reduce both association and dissociation rates. Another possible reason for the lower dissociation rate in the stroma-rich tumor regions was the relaxing of the extracellular matrix, which would prevent the antibody from easily drifting away from the binding zone, thereby resulting in a high fraction of rebinding^11^. The slow dissociation rate resulted in the accumulation of residual antibodies in the stroma-rich tumor areas, even when the systemic antibodies had been largely eliminated, consistent with our immunofluorescence staining results.

We developed two spatially resolved computational models by assuming either heterogeneous binding or heterogeneous distribution between the stroma-rich and stroma-poor tumor regions. We then used competition studies to test which of the two models would consistently predict the observed the dynamic features of cetuximab–EGFR binding. The performance was much better for the model with the heterogeneous binding assumption than with the heterogeneous distribution, indicating that an uneven distribution of antibodies in two tumor areas was not the primary reason for the biphasic declining features in the tumor RO data.

One clarification should be made, namely that the inconsistent predictions produced by the heterogeneous distribution model only suggest that the uneven distributions of antibodies in tumors do not sufficiently explain the observed binding dynamic features. Therefore, this precludes making the implication of even distribution of antibodies in the tumor. One limitation of this study was that the model was developed based on the imaging data in xenografts, which may not recapitulate the complexity of clinical tumors. In addition, the IF staining images were acquired at the end of the BRET imaging study, which was preferably conducted in a longitudinal manner to match with our model simulation.

Overall, in the present study, we combined the strengths of BRET imaging and spatially resolved computational models to evaluate the dynamics of binding of an antibody to its target in living tumors. We demonstrated that spatial heterogeneity exists in antibody-binding profiles between stroma-rich and stroma-poor tumor regions. These findings improve our understanding of the complex antibody targeting process and should aid in the design of antibodies that show more favorable targeting properties.

