## Supplementary Material for "Modeling the dynamics of antibody–target binding in living tumors"

### Supplementary methods

#### Model differential equations

*Plasma*

$$\frac{d}{dt}A_{plasma} = CL_D \cdot C_{peripheral} - \ CL_D \cdot C_{plasma} - CL_p \cdot C_{plasma}$$

*Other tissues*

$$\frac{d}{dt}A_{peripheral} = CL_D \cdot C_{plasma} - \ CL_D \cdot C_{peripheral}$$

*Free cetuximab concentration ($C_1$) in stroma-poor regions*

$$\frac{d}{dt}C_1 = \frac{(1-\sigma_v)\cdot L_p \cdot C_{plasma}}{V_{p\_a}} - \frac{(1-\sigma_L)\cdot L_p \cdot C_1}{V_{p\_a}} - k_{on}\cdot C_1 \cdot R_1 + k_{off\_p}\cdot AR_1$$

*Free EGFR concentration ($R_1$) in stroma-poor regions*

$$\frac{d}{dt}R_1 = k_{syn} - k_{deg}\cdot R_1 - k_{on}\cdot C_1 \cdot R_1 + k_{off\_p}\cdot AR_1$$

*Cetuximab-EGFR complex concentration ($AR_1$) in stroma-poor regions*

$$\frac{d}{dt}AR_1 = k_{on}\cdot C_1 \cdot R_1 + k_{off\_p}\cdot AR_1 - k_{int}\cdot AR_1$$

*Free cetuximab concentration ($C_2$) in stroma-rich regions*

$$\frac{d}{dt}C_2 = \frac{(1-\sigma_v)\cdot L_r \cdot C_{plasma}}{V_{r\_a}} - \frac{(1-\sigma_L)\cdot L_r \cdot C_2}{V_{r\_a}} - k_{on}\cdot C_2 \cdot R_2 + k_{off\_r}\cdot AR_2$$

*Free EGFR concentration ($R_2$) in stroma-rich regions*

$$\frac{d}{dt}R_2 = k_{syn} - k_{deg}\cdot R_2 - k_{on}\cdot C_2 \cdot R_2 + k_{off\_r}\cdot AR_2$$

*Cetuximab-EGFR complex concentration ($AR_2$) in stroma-rich regions*

$$\frac{d}{dt}AR_2 = k_{on}\cdot C_2 \cdot R_2 + k_{off\_r}\cdot AR_2 - k_{int}\cdot AR_2$$

The RO was calculated by the equation below:

$$\mathrm{RO}(\%) = 100 \cdot \frac{AR_1 \cdot V_{p\_a} + AR_2 \cdot V_{r\_a}}{(AR_1 + R_1)\cdot V_{p\_a} + (AR_2 + R_2)\cdot V_{r\_a} + R0 \cdot (\frac{V_{p\_a}}{fav} + \frac{V_{r\_a}}{fav} - V_{p_a} - V_{r_a})}$$

**Supplementary Figure 1**

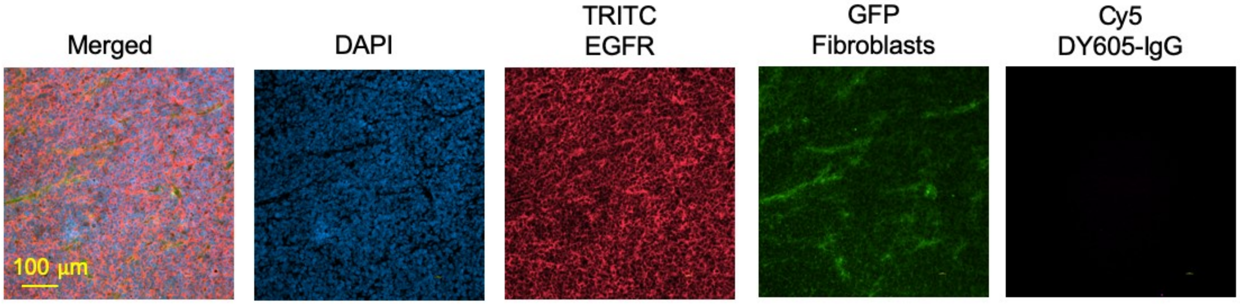


**Figure S1. The Cy5 signals denoted bound DY605-labeled cetuximab rather than non-specific binding.** No residual antibodies were observed for nonspecific IgG, suggesting that the residual antibodies were associated with Fab binding and not with non-specific binding.

**Supplementary Table 1**

*Table S1 Heterogeneous distribution model (TDM) parameter estimations*

| Parameter | Unit | Definition | Estimation (CV%) |
|---|---|---|---|
| $k_{deg}$ | $hr^{-1}$ | EGFR degradation rate | 0.00082 (101%) |
| R0 | nM | EGFR initial concentration in tumor stroma-rich and stroma-poor areas | 0.0027 (292%) |
| $k_{on}$ | $nM^{-1}\cdot hr^{-1}$ | Cetuximab-EGFR association rate | 0.76 (54%) |
| $k_{off}$ | $hr^{-1}$ | Cetuximab-EGFR dissociation rate | 0.57 (20%) |
| $f_t$ | | Ratio of tumor stroma-poor space volume over total space | 0.55 (6.0%) |
| $TBF_p$ | $hr^{-1}$ | Tumor Blood Flow per 1L tumor at stroma-poor regions | 1.02 (37%) |
| $TBF_r$ | $hr^{-1}$ | Tumor Blood Flow per 1L tumor at stroma-rich regions | 0.0067 (39%) |
| $k_{int}$ | $hr^{-1}$ | Cetuximab-EGFR internalization rate | 0.04 (13%) |
